# Molecular tuning of the axonal mitochondrial Ca^2+^ uniporter ensures metabolic flexibility of neurotransmission

**DOI:** 10.1101/685859

**Authors:** Ghazaleh Ashrafi, Jaime de Juan-Sanz, Ryan J. Farrell, Timothy A. Ryan

**Affiliations:** Department of Biochemistry, Weill Cornell Medicine. New York, NY 10065; Sorbonne Universités and Institut du Cerveau et de la Moelle Epinière (ICM) - Hôpital Pitié-Salpêtrière, Inserm, CNRS, Paris, France; The David Rockefeller Graduate Program, Rockefeller University, New York, NY 10065

## Abstract

The brain is a vulnerable metabolic organ and must adapt to different fuel conditions to sustain function. Nerve terminals are a locus of this vulnerability but how they regulate ATP synthesis as fuel conditions vary is unknown. We show that synapses can switch from glycolytic to oxidative metabolism, but to do so, they rely on activity-driven presynaptic mitochondrial Ca^2+^ uptake to accelerate ATP production. We demonstrate that while in non-neuronal cells mitochondrial Ca^2+^ uptake requires elevated extramitochondrial Ca^2+^, axonal mitochondria readily take up Ca^2+^ in response to small changes in Ca^2+^. We identified the brain-specific protein MICU3 as a critical driver of this tuning of Ca^2+^ sensitivity. Ablation of MICU3 renders axonal mitochondria similar to non-neuronal mitochondria, prevents acceleration of local ATP synthesis, and impairs presynaptic function under oxidative conditions. Thus, presynaptic mitochondria rely on MICU3 to facilitate mitochondrial Ca^2+^ uptake during activity and achieve metabolic flexibility.

- Synapses rely on activity-driven mitochondrial ATP synthesis with oxidative fuels
- Mitochondrial Ca^2+^ uptake is required to stimulate ATP synthesis in axons
- The mitochondria Ca^2+^ uptake threshold is lower in axons than in non-neuronal cells
- MICU3 controls the Ca^2+^ sensitivity of MCU in axonal mitochondria

---

In humans, when blood glucose drops below ~ 2.8 mM, severe neurological consequences ensue including delirium and coma (Plum and Posner, 2015). This acute sensitivity to fuel availability is likely exaggerated in the brain compared to other tissues as neurons do not store significant reserves of high-energy molecules and the brain does not appear to carry out beta-oxidation (Schönfeld and Reiser, 2013). In addition, the concentration of glucose in the cerebrospinal fluid is considerably lower (~1mM) than that of blood (~5mM when fasting) (McNay and Gold, 1999).There is accumulating evidence that the brain is capable of adapting to different fuel sources (Vannucci and Vannucci, 2000). For example, a restrictive ketogenic diet is the main therapeutic approach for the treatment of drug-resistant childhood epilepsies (Neal et al., 2008) and chronic blockade of mitochondrial pyruvate uptake makes neurons more reliant on glutamate as a driver of the TCA cycle (Divakaruni et al., 2017). However, the dearth of locally stored biochemical intermediates that can be readily accessed to fuel the various neurophysiological processes implies that ATP must be synthesized on demand regardless of the fuel source. This coupling of activity and ATP production is particularly constrained at nerve terminals, as they typically operate hundreds of microns, or even tens of centimeters away from the cell soma. In support of this view, we previously showed that presynaptic terminals must locally synthesize ATP during activity to meet increased energetic demands (Ashrafi et al., 2017; Rangaraju et al., 2014). Although the brain generally relies exclusively on the delivery of glucose from the body’s other organs to sustain function, the extent to which neurons and synapses rely on glycolytic versus oxidative production of ATP has long been debated (Yellen, 2018). The glycolytic machinery is present in nerve terminals, as 5 of the 10 enzymes required for glycolysis are enriched on synaptic vesicles (Ikemoto et al., 2003; Knull and Fillmore, 1985) including those that produce ATP. We recently showed that glycolysis in nerve terminals can be upregulated in response to activity by AMP-kinase driven insertion of Glut4 glucose transporters into the presynaptic plasma membrane providing an important link connecting supply and demand for ATP at nerve terminals. Whether the glycolytic production of ATP is enough to sustain function will depend on the local glycolytic capacity, which can be limited by the abundance of glycolytic enzymes, the availability of glucose as substrate or both. If the glycolytic capacity is limited, then pyruvate produced either by local glycolysis or from lactate supplied by extra-neuronal sources, such as astrocytes (Pellerin et al., 1998), could in principle fuel local mitochondria to meet ATP demands. However, how synapses might upregulate ATP production under more oxidative conditions remains unknown, (i.e. how can synapses maintain metabolic flexibility while preserving activity-dependent ATP production? To answer this fundamental question, we examined the regulation of local ATP production during electrical activity in nerve terminals when the carbon source for ATP generation was switched from extracellular glucose to extracellular lactate and pyruvate. We show that rather than a feedback mechanism such as the one used to upregulate glycolysis, under oxidative conditions, nerve terminals rely on a feed-forward mechanism via mitochondrial Ca^2+^ uptake to accelerate ATP production. Classic experiments using a first-generation genetically-encoded and mitochondrially targeted Ca^2+^ indicator (Rizzuto et al., 1993) demonstrated that Ca^2+^ uptake into mitochondria likely occurs only at sites of close apposition between an intracellular Ca^2+^ store (e.g. the endoplasmic reticulum) and mitochondria, consistent with in-vitro data demonstrating that mitochondria do not readily take up Ca^2+^ unless exposed to high Ca^2+^ concentrations (Sparagna et al., 1994). We show that presynaptic mitochondria, in contrast to those in non-neuronal cells, have a much lower threshold for Ca^2+^ uptake, and that this property is conveyed by MICU3, a brain-specific regulator of the mitochondrial Ca^2+^ uniporter (MCU), freeing neurons from the constraint of relying on an intracellular Ca^2+^ store for mitochondrial Ca^2+^ uptake. In the absence of either MICU3 or MCU, the metabolic plasticity of presynaptic terminals is diminished, resulting in impairment of synaptic function.

## Results

### Upregulation of mitochondrial ATP synthesis during electrical activity at nerve terminals

We previously demonstrated that electrical-activity-driven upregulation of nerve terminal ATP synthesis is required to sustain synaptic function as the energy needs of vesicle recycling rapidly outstrips the availability of local ATP (Ashrafi et al., 2017; Rangaraju et al., 2014). Subsequent experiments showed that one mechanism by which ATP synthesis can be upregulated is by increasing glucose uptake (Ashrafi et al., 2017), thereby increasing glycolytic flux. These experiments imply that under these conditions, production of the primary fuel for oxidative phosphorylation, pyruvate, would also accelerate. Whether mitochondria upregulate ATP synthesis simply due to an increase in substrate, or because biochemical turnover in the mitochondrial matrix is being accelerated is unknown. To address this issue, we monitored ATP levels in nerve terminals of dissociated primary hippocampal neurons using SYN-ATP, a genetically-encoded and synaptically-targeted ATP indicator (Rangaraju et al., 2014), where the normal extracellular fuel source, glucose, has been completely replaced with a mixture of lactate and pyruvate to force neuronal reliance on mitochondrial oxidative metabolism. Robust AP-firing (600 AP at 10 Hz) under these conditions however did not impact ATP levels (Fig 1A and B). This finding implies that either such level of electrical activity does not consume substantial amounts of ATP, or that despite the increase in ATP consumption, ATP production is accelerated to match these demands. Our previous work (Rangaraju et al., 2014) showed that this level of AP firing leads to significant ATP consumption. Here under elevated lactate/pyruvate it is possible that we have simply masked the ATP consumption by increasing basal ATP production levels. However, blocking mitochondrial ATP production using the F_1_-F_0_ ATP synthase inhibitor oligomycin, leads to an abrupt drop in ATP levels during AP firing (Fig. 1A and B). These experiments show that, at nerve terminals, mitochondria respond to electrical activity by upregulating ATP production to precisely match the increased demand for energy. In order to determine if this acceleration in mitochondrial ATP synthesis is important to sustain nerve terminal function, we monitored AP-driven synaptic vesicles (SV) exocytosis, endocytosis and reacidification using vGLUT1-pHluorin (vGLUT1-pH) (Voglmaier et al., 2006). Replacing extracellular glucose with a mixture of lactate and pyruvate has no impact on the kinetics of SV recycling (Fig.1C), demonstrating that synapses have a profound metabolic flexibility with respect to the fuel source. Acute blockade of mitochondrial ATP production however leads to a complete arrest of SV recycling (Fig. 1D and E), as expected, since nerve terminals completely rely on oxidative fuel production under these conditions. Thus, as with glycolytic conditions, nerve terminals also upregulate ATP synthesis under oxidative conditions, but in this case by accelerating mitochondrial function.

**Figure 1.**
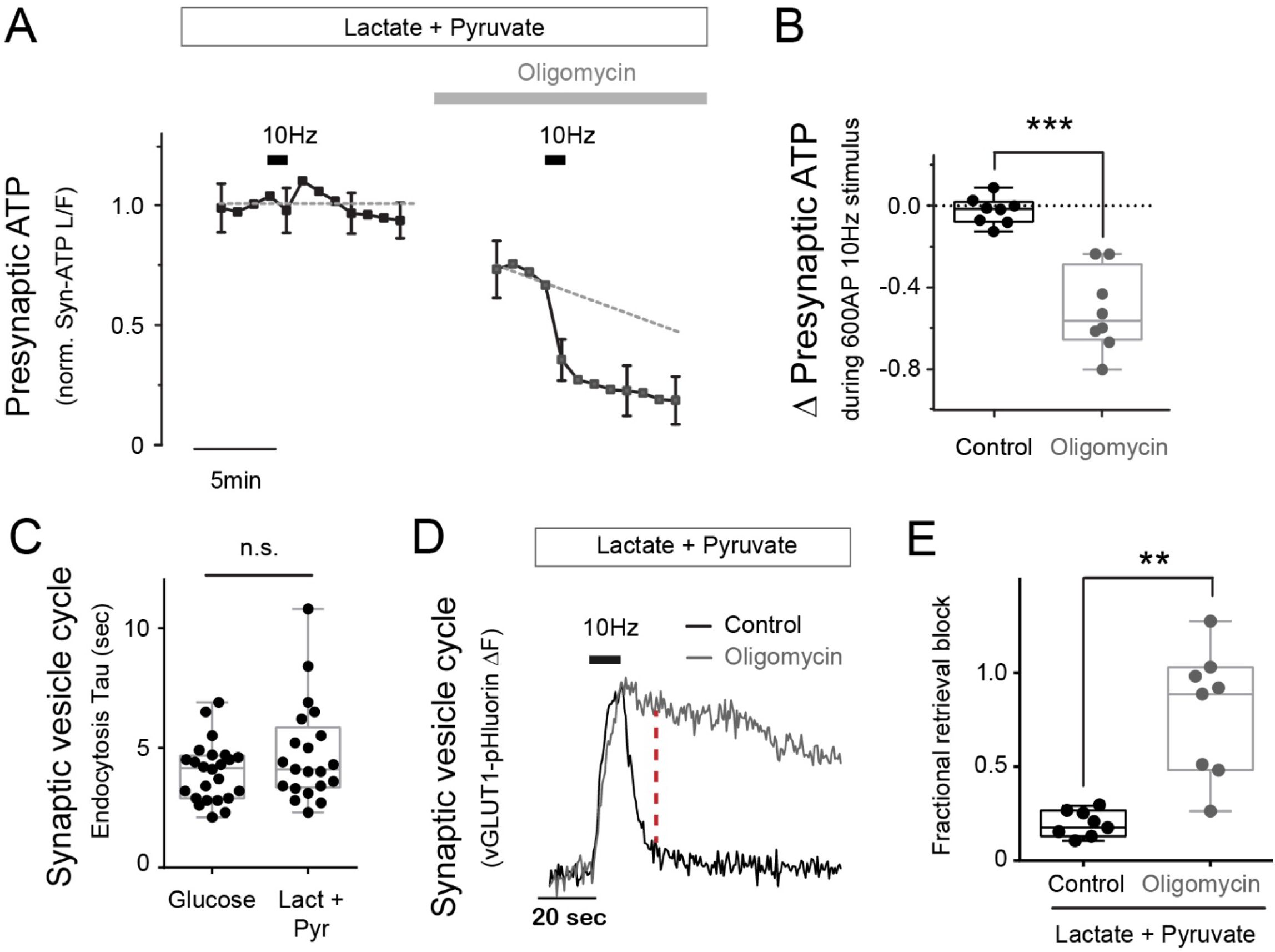
Activity-driven mitochondrial ATP production in nerve terminals. (A) Average normalized traces of presynaptic ATP, measured with the luminescent reporter Syn-ATP. Neurons were electrically stimulated with 600AP at 10Hz (black bar) before and after a 5-minute treatment with oligomycin in the presence of lactate and pyruvate but with no glucose. The grey dashed line represents linear fitting to the points prior to stimulation. (B) Change in normalized ATP levels immediately after stimulation (n= 8 cells). Average change in ATP: control, −0.02 ± 0.02; oligomycin, −0.51 ± 0.07. (C) Endocytosis time constants (seconds) of vGLUT1-pH in neurons stimulated in the presence of glucose or lactate and pyruvate. n= 21-24 cells. Mean (sec); glucose, 4.0; lact+pyr, 4.8. (D) Sample vGLUT1-pH traces in response to 100AP, 10Hz (black bar) before and 5 minutes after oligomycin treatment in the presence of lactate and pyruvate. (E) Fractional retrieval was calculated as the fraction of vGLUT1-pH signal remaining after 2 times the endocytic constant of control (red dashed line in D) after stimulation (0 and 1 denote complete and no endocytosis, respectively). See STAR Methods. n = 8 cells. Mean; control, 0.2; oligomycin 0.8. Error bars are SEM. **p<0.01, ***p<0.001. n.s. not significant. The box-whisker plot represents median (line), 25th–75th percentile (box), and min–max (whisker).

### MCU is required for presynaptic mitochondrial uptake and acceleration of ATP production under oxidative conditions

It has long been hypothesized that mitochondrial ATP production is accelerated by Ca^2+^ uptake into mitochondria (Heineman and Balaban, 1990), although detailed tests of this hypothesis in intact cells has been lacking. We took advantage of the identification of the principal molecular pathway responsible for mitochondrial Ca^2+^ uptake, the mitochondrial Ca^2+^ uniporter (MCU) (Baughman et al., 2011; De Stefani et al., 2011), to see if presynaptic mitochondrial Ca^2+^ uptake during activity is necessary to accelerate ATP production. We monitored mitochondrial Ca^2+^ uptake in axons during AP firing (Fig. 2A) using an improved mitochondrial-targeting of GCaMP6f (mito^4x^-GCaMP6f, see methods). As all GCaMPs are also sensitive to pH we verified that the stimulus levels used did not impact mitochondrial pH using mito^4x^-pHluorin (Fig. S1A). Even brief bursts of AP firing leads to very robust Ca^2+^ uptake in axonal mitochondria (Fig. 2A and B). This uptake was heavily suppressed when expression of MCU was transiently ablated using an shRNA targeting MCU (Fig. 2A and B), as validated by qPCR (Fig. S1C). This finding is consistent with previous work conclusively demonstrating that MCU is the primary rapid uptake pathway for Ca^2+^ in mitochondria (Pan et al., 2013). MCU ablation, however, did not significantly impact resting mitochondrial matrix Ca^2+^ levels (Fig. S1D). We found that neurons do not tolerate overexpression of MCU, consistent with previous reports in that over-expression of MCU increases excitotoxicity in neurons (Qiu et al., 2013). Similarly, when attempting to rescue MCU ablation with an shRNA-insensitive variant of MCU few neurons survived. Those that did had partially restored mitochondrial Ca^2+^ uptake (Fig. S1B).

**Figure 2.**
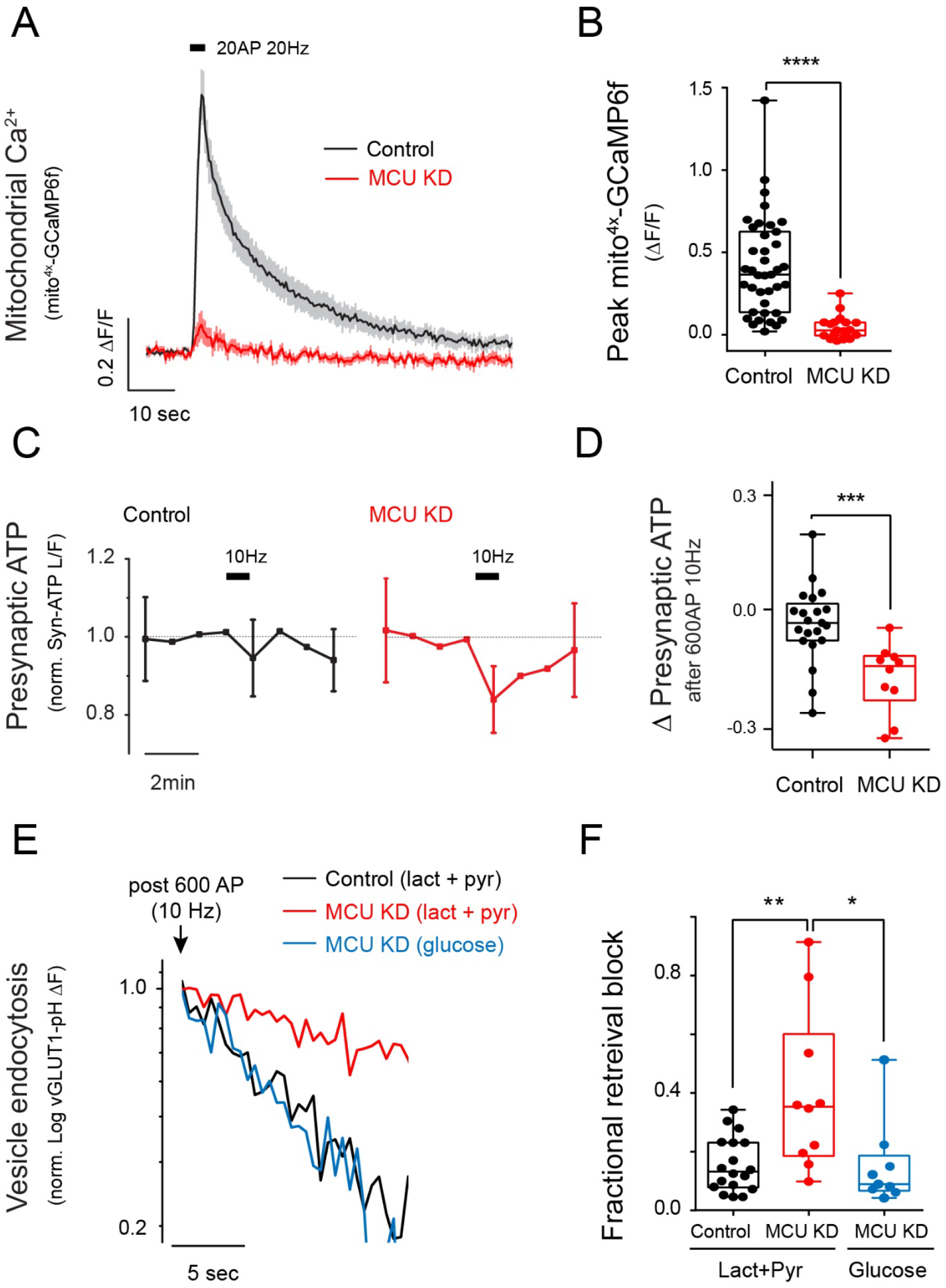
MCU is required for mitochondrial Ca^2+^ uptake and activity-driven ATP production. (A, B) Average traces of Mito^4x^-GCaMP6 (A) showing mitochondrial Ca^2+^ uptake in control and MCU KD axons stimulated with 20AP at 20Hz. (B) Peak responses of Mito^4x^-GCaMP6 (ΔF/F) following stimulation. n = 19-39 cells. mean (ΔF/F); control, 0.41; MCU KD, 0.04. (C, D) Normalized presynaptic ATP traces (C) in control and MCU KD neurons stimulated with 600AP at 10Hz. The grey dashed line represents linear fitting to the points prior to stimulation. (D) Change in normalized ATP levels immediately after stimulation. n= 9-21 cells. Mean; control, −0.04; MCU KD, −0.17. (E, F) Sample semi-log plot of vGLUT1-pH traces (E) following stimulation with 600AP at 10Hz in control (black) and MCU KD neurons (red) supplied with lactate and pyruvate. Blue trace shows the response of the same MCU KD neuron shown in red but in the presence of media containing glucose. (F) Fractional retrieval block calculated as described in STAR methods. n = 9-18 cells. Mean; control, 0.16; MCU KD (lact+pyr), 0.40; MCU KD (glucose), 0.15. Unless otherwise indicated, all experiments were performed in the presence of lactate and pyruvate. Error bars are SEM. *p<0.05, **p<0.01 ****p<0.0001.

Many lines of evidence support the idea that mitochondrial Ca^2+^ uptake can accelerate the tricarboxylic acid cycle (TCA) to enhance ATP production (Glancy and Balaban, 2012). Since ablation of MCU severely impairs mitochondrial uptake, we measured ATP levels at synapses during 1 min of AP firing in neurons that lack MCU. Unlike control neurons, where ATP levels are unperturbed by activity, in the absence of MCU, ATP levels declined during the period of activity and slowly recovered (Fig. 2C and D). Loss of MCU, however, is not equivalent to blocking the F_1_-F_0_ ATP synthase. These experiments demonstrate that axonal mitochondrial Ca^2+^ uptake is specifically needed to stimulate ATP synthesis during robust neuronal activity. In order to determine if the inability to accelerate ATP synthesis during activity impacts nerve terminal function, we measured the kinetics of SV recycling using vGLUT1-pH as above. These experiments show that in neurons that lack MCU, SV endocytosis and reacidification are slowed (Fig. 2E and F). These experiments are carried out under purely oxidative conditions (lactate/pyruvate, zero glucose) where all new ATP synthesis depends on oxidative phosphorylation. In principle, synaptic dysfunction in the absence of MCU might have arisen from downstream consequences of chronic mitochondrial dysfunction. However, simply replacing the fuel source with glucose completely restored synaptic function even in the absence of MCU (Fig. 2E and F). Thus, loss of MCU-mediated Ca^2+^ uptake specifically impairs synapse function under oxidative, not glycolytic conditions, implying the defects are not pleiotropic in nature. These experiments indicate that MCU is required for the ability of nerve terminals to effectively upregulate mitochondrial ATP production in switching from glycolytic to oxidative conditions.

### Presynaptic mitochondria do not rely on the ER as a source of Ca^2+^ uptake

Mitochondrial Ca^2+^ uptake is activated by Ca^2+^ but is thought to be highly constrained requiring Ca^2+^ levels that would only be satisfied when mitochondria are in close apposition to a Ca^2+^ release site, such as IP_3_ receptors or Ryanodine receptors, on the endoplasmic or sarcoplasmic reticulum. This idea is supported by the fact that isolated mitochondria only take up Ca^2+^ when perfused with high Ca^2+^ concentrations (Gunter et al., 1994), and the observation that perfusion of permeabilized cells with IP_3_ triggers mitochondrial Ca^2+^ uptake, while perfusion of sub-micromolar Ca^2+^ does not (Rizzuto et al., 1993). This cooperation between mitochondria and the ER would therefore implicate the latter organelle in playing a critical role in controlling Ca^2+^-mediated regulation of oxidative phosphorylation. In order to determine if the ER plays such a role in activity-driven mitochondrial Ca^2+^ uptake in axons, we examined mitochondrial Ca^2+^ uptake before and after blocking ER Ca^2+^ handling (Fig. 3C and D). During AP firing, axonal ER in these neurons acts as a net sink for Ca^2+^ (Fig 3A), as we previously demonstrated using a new genetically-encoded and ER-targeted ER Ca^2+^ sensor based on GCaMP6 but with an affinity altered to match more closely the resting ER Ca^2+^level (de Juan-Sanz et al., 2017). Inhibition of the sarcoplasmic/endoplasmic reticulum Ca^2+^ ATPase (SERCA) with cyclopiazonic acid (CPA) eliminates ER Ca^2+^ signaling (Fig. 3A and B) because in the absence of SERCA activity, Ca^2+^ leaks out of the ER. We previously showed that following SERCA block, ER Ca^2+^ decreases with a time constant of < 1 min under these conditions (de Juan-Sanz et al., 2017). Mitochondrial Ca^2+^ uptake, however, was unaffected by SERCA blockade when probed several minutes after CPA addition (Fig. 3C and D) indicating that activity-driven mitochondrial Ca^2+^ uptake does not depend on the ER to satisfy potential constraints in Ca^2+^-dependent MCU activation.

**Figure 3.**
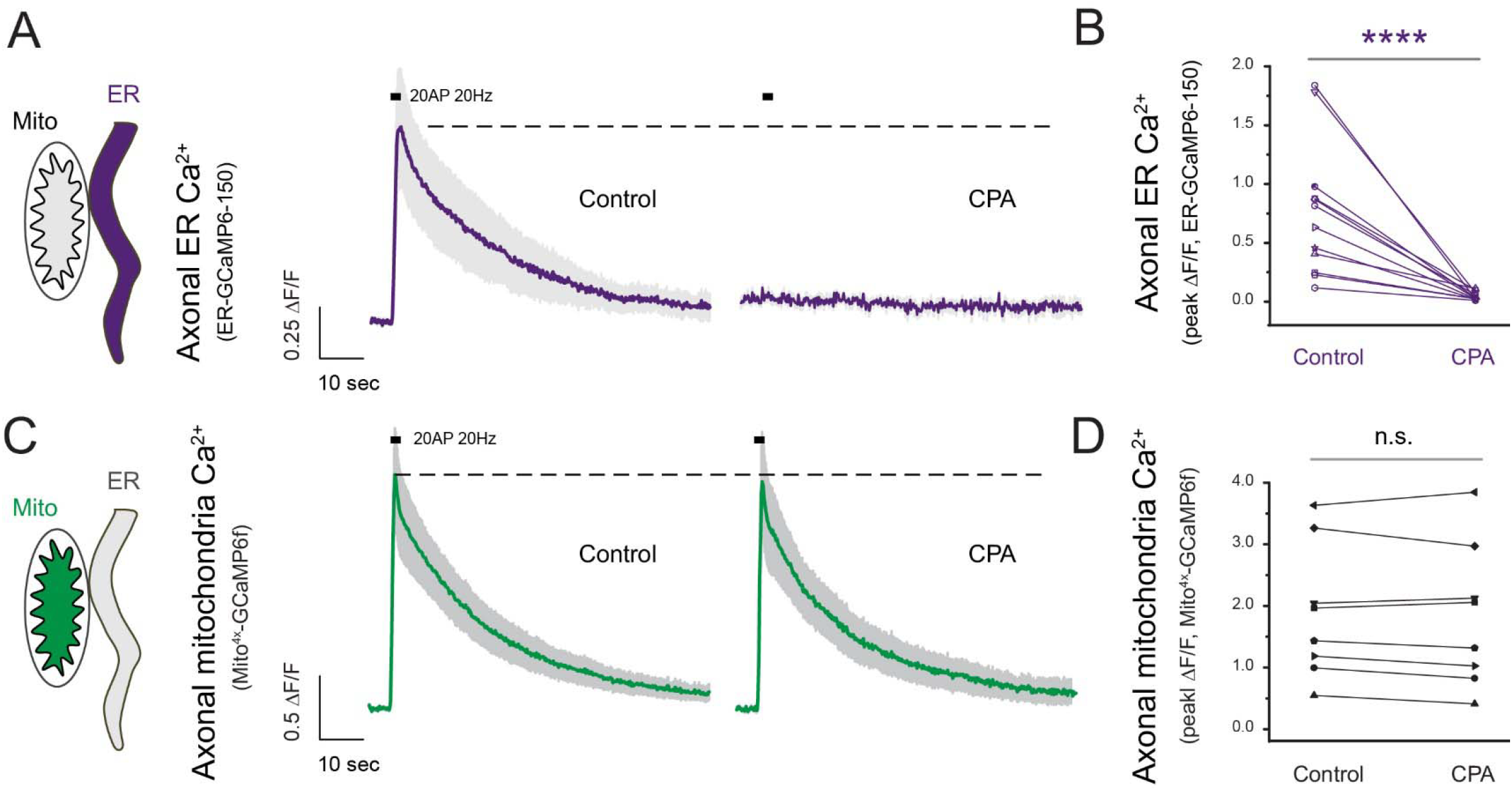
Axonal mitochondria do not rely on the ER for Ca^2+^ uptake. (A-D) Average traces of ER-GCaMP6-150 (A), or Mito^4x^-GCaMP6 (C) in control and neurons treated with the SERCA inhibitor CPA to silence axonal ER Ca^2+^ responses (A, B). Paired comparison of peak ER-GCaMP6-150 (B) or mito^4x^-GCaMP6 (D) ΔF/F responses before and after CPA treatment. n = 8-12 cells. Mean peak ER Ca^2+^ response (ΔF/F); control, 0.77; CPA-treated, 0.04. Mean peak mitochondrial Ca^2+^ response (ΔF/F); control, 1.88; CPA-treated, 1.82. Experiments were performed in the presence of glucose. Error bars in traces are SEM.

### Higher Ca^2+^ concentrations are needed to trigger mitochondrial Ca^2+^ uptake in non-neuronal cells compared to nerve terminals

The finding that axonal mitochondria are freed from the constraints of relying on ER Ca^2+^ to activate MCU implies that either axonal mitochondria normally occupy a privileged location close to sites of plasma-membrane Ca^2+^ entry, or that the machinery associated with MCU activation is specifically tuned to operate with lower Ca^2+^ concentrations in axons. To distinguish between these two scenarios, we developed a novel approach that allowed us to compare the Ca^2+^ signals needed on the outer mitochondrial membrane (OMM) to trigger Ca^2+^ uptake into the mitochondrial matrix in a fibroblast versus a neuronal axon. To measure OMM Ca^2+^ signals, we fused the genetically-encoded red-shifted Ca^2+^ indicator jRCaMP1b (Dana et al., 2016) to a portion of the OMM protein TOM20 and expressed this construct in neurons. Ca^2+^ influx in axons can be robustly stimulated and controlled by varying AP firing at a fixed extracellular Ca^2+^ concentration. Over many experiments across axons of many individual neurons, we compared the mitochondrial matrix GCaMP6f signal for a given stimulus condition with the OMM jRCaMP1b signal for the same stimulus condition. We then correlated the matrix Ca^2+^ signal obtained for a given stimulus with the OMM signal for the same stimulus. These experiments show that axonal mitochondrial Ca^2+^ uptake appears non-linear. With minimal Ca^2+^ entry (e.g. a 3 AP stimulus) which generates a robust OMM signal, the matrix signal is negligible (Fig. 4A).

**Figure 4.**
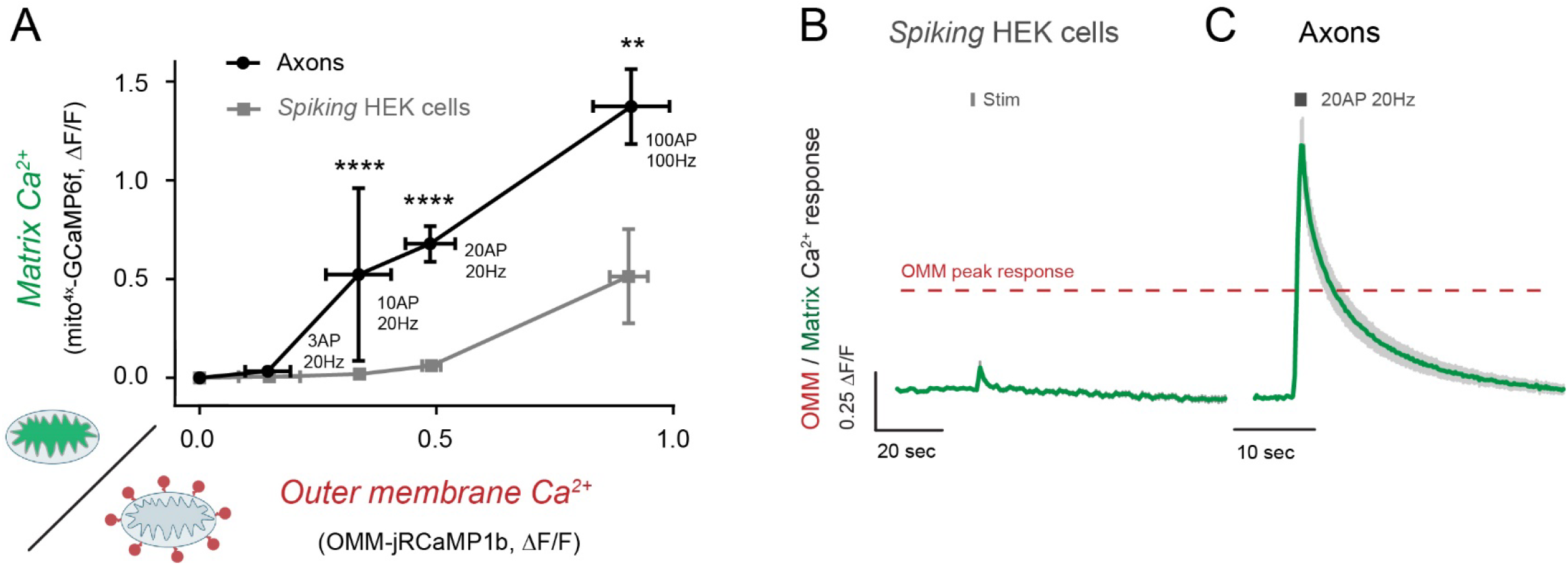
Lower Ca^2+^ levels are needed to trigger mitochondrial Ca^2+^ uptake in nerve terminals than in non-neuronal cells. (A-C) Correlation of the mitochondrial matrix-targeted mito^4x^-GCaMP6 and the outer membrane-targeted OMM-jRCaMP1b signal (A) in response to a given stimulus in mitochondria of spiking HEK cells and neuronal axons. Responses in spiking HEK cells were binned across OMM responses for ease of comparison with f neurons. (B, C) Averaged traces of matrix (green) Ca^2+^ fluxes in spiking HEK cells (B) and neuronal axons (C) in response to 20AP at 20Hz in neurons and the equivalent Ca^2+^ increase in the outer membrane (OMM) of HEK cells, measured by quantifying OMM Ca^2+^ signals (red) in both cell types (traces not shown for clarity, but average OMM peak response is denoted with a red dashed line). Experiments were performed in the presence of glucose. Error bars are SEM.

However, increasing the OMM signal by ~ 2.5 fold led to a greater than 10-fold increase in the matrix signal (Fig. 4A and C). In order to determine the sensitivity of non-neuronal mitochondria we developed a novel approach to allow us to reproducibly induce transient changes in cytosolic Ca^2+^ that do not depend on ER function in human embryonic kidney (HEK) cells. We made use of the development of “spiking” HEK cells (Park et al., 2013), that constitutively express Na_v_1.3 and K_IR_2.1 which allow these cells to fire an AP upon brief extracellular field stimulation (Fig. S2). We transiently transfected these cells with our mitochondrial OMM and matrix reporters (see STAR Methods) along with plasmids encoding the three genes necessary for the surface expression of a Cav2.1 voltage-gated Ca^2+^ channel (α1b, α2δ1 and β1). A single AP stimulus in these cells leads to a very robust and transient influx of Ca^2+^ across the plasma membrane (Fig. S2). By varying the concentration of extracellular Ca^2+^, we could vary the size of the OMM signal and determine the correlation of mitochondrial Ca^2+^ uptake with OMM Ca^2+^ in these non-neuronal cells. These experiments showed that unlike neurons that showed robust mitochondrial Ca^2+^ uptake for relatively small changes in OMM Ca^2+^, mitochondria in HEK cells remained unresponsive over a much wider range OMM Ca^2+^ signals (Fig. 4 A and B). The transition from minimal Ca^2+^ uptake to robust Ca^2+^ uptake appeared, as with neuronal mitochondria, to be quite non-linear where a doubling of OMM Ca^2+^ signal (from a ΔF/F of 0.42 to 0.84) led to a greater than 10-fold increase in the matrix Ca^2+^ signal. However, the apparent set point for this transition was ~3 fold higher in fibroblasts compared to neurons. These experiments demonstrate that the likely reason axonal mitochondria do not depend on ER function is that their machinery for mitochondrial Ca^2+^ uptake is tuned to be responsive to smaller OMM Ca^2+^ signals such that Ca^2+^ entry from voltage-gated-Ca^2+^ channels for modest AP firing is sufficient to trigger MCU activation.

### MICU3 shifts the Ca^2+^ sensitivity of presynaptic mitochondrial Ca^2+^ uptake

The experiments above demonstrate the axonal mitochondria have a significantly different Ca^2+^ sensitivity for activating mitochondrial Ca^2+^ uptake than non-neuronal mitochondria. The Ca^2+^-dependent activation of MCU is conferred by a set of EF-hand containing regulatory proteins, MICUs, that assemble around MCU in the mitochondrial intramembrane space (Marchi and Pinton, 2014). In most tissues, this is conveyed by MICU1 and MICU2 where it is thought that MICU1 is responsible for keeping MCU in a closed state (Mallilankaraman et al., 2012) and MICU2 allows Ca^2+^-dependent MCU opening (Patron et al., 2014). MICU3 is a brain-specific isoform that is exclusively expressed in neurons (Plovanich et al., 2013) and was recently shown to enhance mitochondrial Ca^2+^ uptake when expressed in non-neuronal cells (Patron et al., 2019). We reasoned that MICU3 might be responsible for the lower threshold of Ca^2+^-driven MCU activated in axonal mitochondria. To test this idea, we used shRNA-mediated ablation of MICU3 (Fig. S3B) and examined axonal mitochondrial Ca^2+^ uptake in dissociated primary hippocampal neurons using mito^4x^-GCaMP6f. For our standard conditions (20 Hz AP firing), loss of MICU3 led to a very significant suppression of mitochondrial Ca^2+^ uptake (Fig. 5A-C), similar to that obtained with ablation of MCU (Fig. 2A and B), which was reversed upon re-expression of an shRNA-insensitive variant of MICU3 (Fig. S3A). However, when we used a more intense stimulus (100 AP at 100 Hz, which is outside the likely physiological range for hippocampal neurons) that leads to greater elevations in cytoplasmic Ca^2+^ (Fig. S3F and G), axonal mitochondria showed robust Ca^2+^ uptake even in the absence of MICU3 (Fig. 5A-C). Although, for milder stimuli, loss of MICU3 suppresses mitochondrial Ca^2+^ uptake to the same extent as loss of MCU, with more intense stimulation of MCU-lacking mitochondria remained unresponsive (Fig. 5C). Thus, without MICU3, axonal mitochondria require higher Ca^2+^ to trigger Ca^2+^ uptake. Measurements of the OMM signal required to activate MCU in axonal mitochondria in the absence of MICU3 indicates that is now like that measured in non-neuronal cells (Fig. S3G). Conversely, expression of MICU3 in spiking HEK cells shifted the mitochondria to a much more Ca^2+^-sensitive state and were now similar to mitochondria in neuronal axons (Fig. 5D-F). Thus, MICU3 expression is sufficient to significantly switch the ability of a mitochondrion to respond to smaller Ca^2+^ elevations.

**Figure 5.**
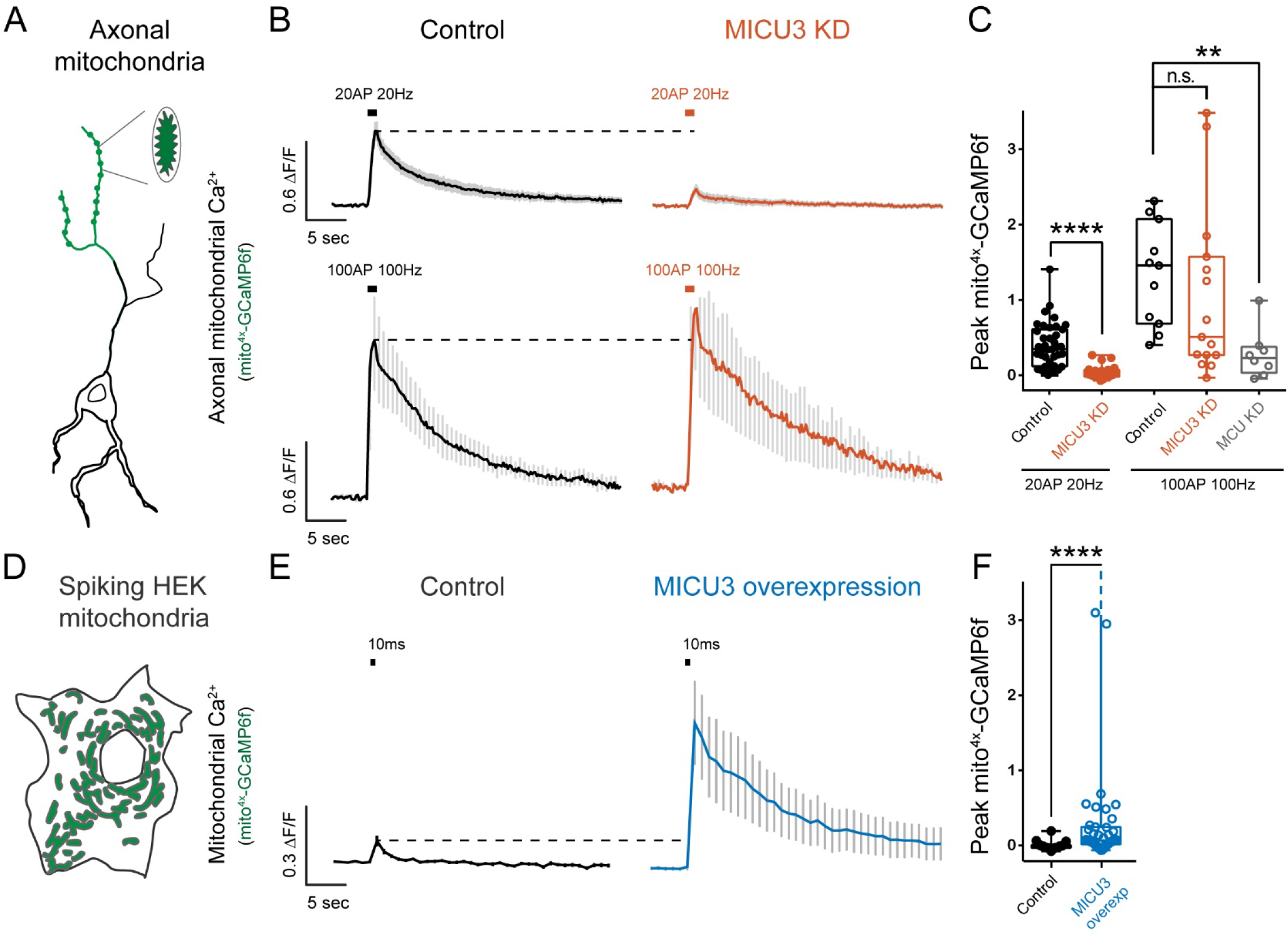
MICU3 shifts the Ca^2+^ sensitivity of presynaptic Ca^2+^ mitochondrial uptake. (A-C) Average traces of mito^4x^-GCaMP6 (B) in neuronal axons (schematic in A) stimulated with 20AP, 20Hz (B, top panel) or 100AP, 100Hz (B, bottom panel) in control and MICU3 KD neurons. (C) Peak mito^4x^-GCaMP6 ΔF/F following stimulation. n = 8-39 cells. mean (ΔF/F); control (20AP, 20Hz) (same as in Fig. 2A and B), 0.41; MICU3 KD (20AP, 20Hz), 0.04; control (100AP, 100Hz), 1.33; MICU3 (100AP, 100Hz), 1.04; MCU KD (100AP, 100 Hz), 0.28. (D-F) Average traces of mito^4x^-GCaMP6 (E) in control and MICU3-overexpressing spiking HEK cells (schematic in D) stimulated for 10 msec. (F) Peak mito^4x^-GCaMP6 ΔF/F following stimulation. n = 34-43 cells. mean; control, 0.03; MICU3 overexpression, 0.4. Error bars are SEM. p>0.05, **p<0.01, ****p<0.0001

### MICU3 is required for feed-forward regulation of ATP production in nerve terminals

We showed that Ca^2+^ uptake in axonal mitochondria is a critical aspect of a feed-forward mediated regulation of ATP synthesis, which under oxidative conditions is in turn critical for presynaptic function (Fig. 2). We therefore examined whether MICU3-enhanced Ca^2+^-sensitivity of Ca^2+^ uptake in neuronal mitochondria contributes to metabolic regulation at nerve terminals. Measurements of ATP at nerve terminals showed that when MICU3 expression is ablated, ATP levels cannot be sustained during 60 s of 10 Hz AP firing (Fig. 6A and B), similar to what was found in neurons lacking MCU. This inability to sustain ATP synthesis resulted in slowing of SV recycling as probed with vGLUT1-pH (Fig. 6C and D). Switching the fuel source to glucose, however, fully restored SV recycling kinetics, indicating, as with loss of MCU, that chronic loss of MICU3 does not lead to pleiotropic impairment of presynaptic machinery. These findings demonstrate that MICU3 is critical in neurons to allow the transition from glycolytic to oxidative metabolism in supporting activity-driven ATP production.

**Figure 6.**
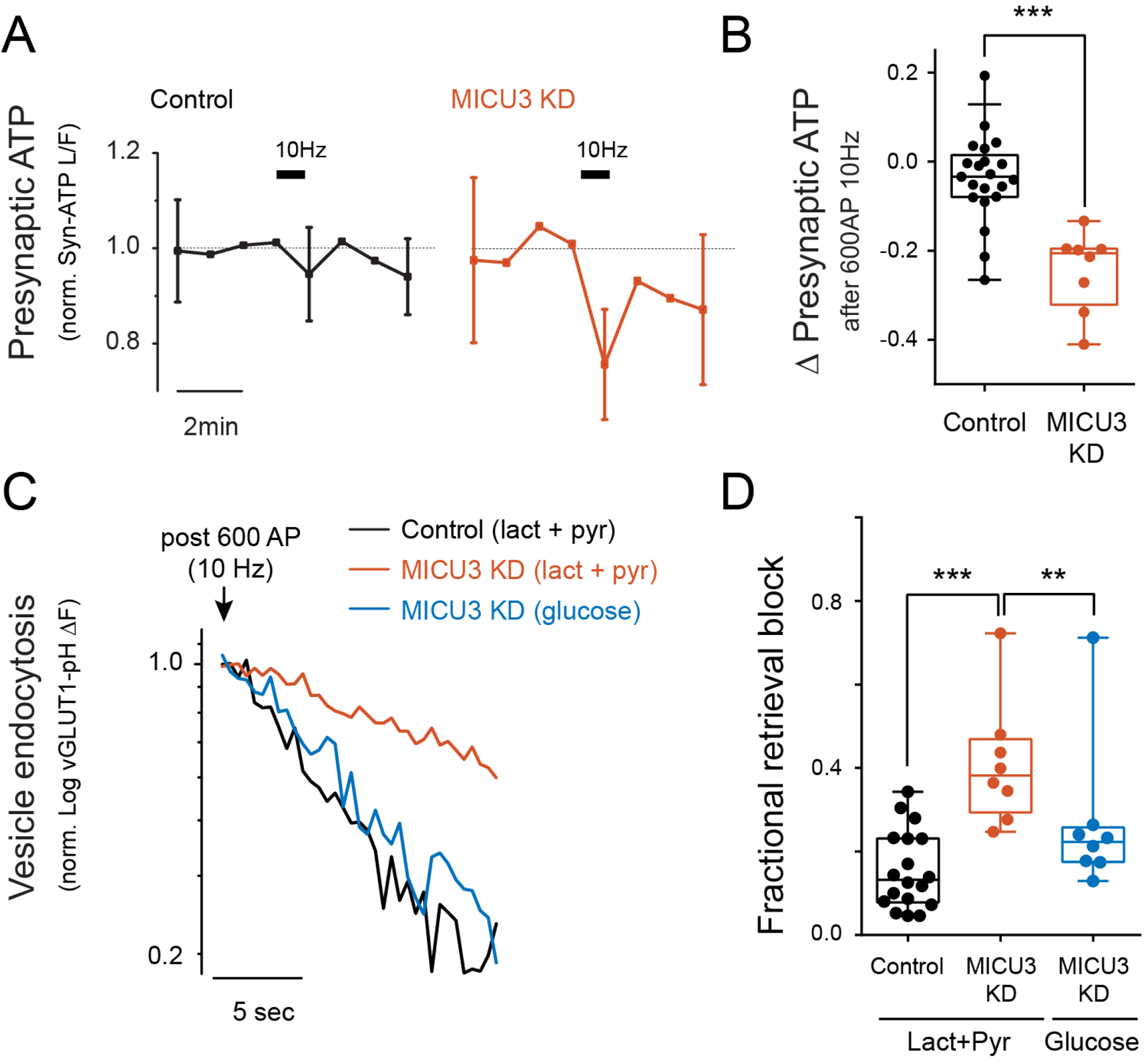
MICU3 is required for feed-forward regulation of ATP production in nerve terminals. (A and B) Normalized presynaptic ATP traces (A) in control and MICU3 KD neurons stimulated with 600AP, 10Hz supplied with lactate and pyruvate. The grey dashed line represents linear fitting to points prior to stimulation. (B) Change in normalized ATP levels immediately after stimulation. n= 8-21 cells. Mean; control (same as in Fig. 2C and D), −0.04; MICU3 KD, −0.25. (C and D). Sample semi-log plot of vGLUT1-pH traces (C) following stimulation with 600AP, 10Hz in control and MICU3 KD neurons supplied with lactate and pyruvate or glucose. (D) Fractional retrieval block as described in STAR Methods: Control (same as in Fig. 2E and F), 0.16; MICU3 KD (lact+pyr), 0.41; MICU3 KD (glucose), 0.27. Error bars are SEM. **p<0.01, ***p<0.001

## Discussion

The metabolic vulnerability of the brain underpins the importance of understanding how fuel is delivered, how brain metabolism is regulated, and the efficiency with which neuronal processes utilize metabolic products including ATP. Inevitably, any disruption in these metabolic processes could lead to neuronal dysfunction and cognitive decline. The processes of SV cycling and maintaining ionic homeostasis during neuronal firing come at steep metabolic costs and require ATP production to adjust according to demand. We have previously shown that ATP levels remain unperturbed at nerve terminals when going from a resting state to constant AP firing despite increased energy consumption (Rangaraju et al., 2014), due to concomitant upregulation of presynaptic ATP production. Although glycolytic ATP synthesis is stimulated during activity (Ashrafi and Ryan, 2017), the ATP produced solely by glycolysis may be insufficient to support function under physiological concentrations of brain glucose, which is significantly lower than the blood (McNay and Gold, 1999). Additionally, it has long been argued that lactate derived from peri-synaptic astrocytes is the main fuel sustaining synapse function (Pellerin et al., 1998). This intrinsic variability in how synapses are fueled implies that they likely upregulate ATP synthesis in a fuel-dependent fashion. Thus, proper regulation of ATP production in response to activity requires distinct mechanisms suited for different metabolic conditions. We show here that nerve terminals are metabolically flexible and can carry out vesicle recycling when dispensing entirely with glycolysis (Fig. 1C), in agreement with previous reports (Pathak et al., 2015). Measurements of ATP concentration during activity under these conditions indicate that, as with glycolysis, mitochondrial ATP production is also upregulated by AP firing. Such metabolic flexibility is notable given the observation that only about 50% of hippocampal *en-passant* boutons contain mitochondria (Kang et al., 2008) and therefore have immediate access to mitochondrially-derived ATP. We previously observed that resting presynaptic ATP levels are sufficient to energetically sustain a 10 AP stimulus, indicating that activity-driven ATP production is only required during repetitive stimuli over a few seconds (Ashrafi et al., 2017). Given that the acceleration of mitochondrial ATP production occurs during repetitive stimuli, we propose that the distance between a nerve terminal and the closest mitochondrion, on average about 3 μm (Smith et al., 2016), is likely to be close enough to allow for newly-synthesized ATP to diffuse and reach the firing terminals, as previously reported (Pathak et al., 2015).

Although mitochondrial Ca^2+^ uptake has long been proposed to modulate the capacity of mitochondria to produce ATP, to our knowledge, the experiments in Fig. 2D are the first to demonstrate that, in intact cells, mitochondrial Ca^2+^ uptake is critical for metabolic support. Here, Ca^2+^ is used as a feed-forward signal serving as a surrogate for the increased energetic demands associated with Ca^2+^ entry. This system compliments but differs from the regulation of glycolysis, where a feedback mechanism, based on AMPK activation (and therefore reflecting the detection of a transient energy deficit) regulates glucose uptake. The downstream mechanisms by which mitochondrial Ca^2+^ uptake drives ATP synthesis have not been explored in neurons.

In non-neuronal cells, mitochondrial Ca^2+^ uptake has been shown to take place where mitochondria form sites of close apposition with the ER (Rizzuto et al., 1993; Rizzuto et al., 1998). This spatial relationship is thought to allow MCU activation to have a high threshold with respect to Ca^2+^ concentration, as upon ER Ca^2+^ release the outer and inner mitochondrial membranes would experience local micro-domains of high Ca^2+^ enabling MCU to open. These organelle-organelle contacts between mitochondria and the ER have been identified in most cell types examined, including neurons (Wu et al., 2017). Presumably, if nerve terminals were to rely upon ER Ca^2+^ release to trigger mitochondrial Ca^2+^ uptake, an initial Ca^2+^ influx through voltage-gated Ca^2+^ channels would trigger Ca^2+^ release from the ER. By silencing axonal ER Ca^2+^ fluxes (de Juan-Sanz et al., 2017), we demonstrated that axonal mitochondrial Ca^2+^ uptake during AP firing is completely independent of the ER’s ability to flux Ca^2+^. Given the long-held view that mitochondrial Ca^2+^ uptake is tightly controlled by its intimate spatial proximity with the ER, two possibilities arose: 1) in axons, mitochondria are positioned very close to voltage-gated Ca^2+^ channels on the plasma membrane such that they always sense high Ca^2+^ during activity or 2) the molecular machinery for mitochondrial Ca^2+^ uptake in axonal mitochondria is different from other tissues. Our quantitative measurement of the cytoplasmic signals that can drive mitochondrial Ca^2+^ uptake (Fig. 4) demonstrated that axonal mitochondria have a lower threshold for triggering mitochondrial Ca^2+^ uptake than mitochondria in non-neuronal cells and therefore do not need to be positioned close to other Ca^2+^ sources. However, the molecular machinery controlling mitochondrial Ca^2+^ uptake appears to be different than in neurons than in other tissues. It is possible that in other subcellular regions of a neuron, mitochondrial Ca^2+^ uptake has a greater dependence on Ca^2+^ release from the ER, as has been reported for mitochondrial Ca^2+^ uptake in dendrites (Hirabayashi et al., 2017). While it has been widely assumed that the physical contacts between the ER and mitochondria facilitate mitochondrial Ca^2+^ uptake, our data imply that these contacts in axons likely serve different roles, for example regulating lipid homeostasis, as the presence of MICU3 relieves the constraints of needing high local Ca^2+^ to drive MCU opening.

It is interesting to note than even across non-neuronal tissues, MCU-based Ca^2+^ uptake activity is highly variable (Fieni et al., 2012). This may be, in part, due to differential tissue expression of the MCU regulators, MICUs. Unlike MICU1 and MICU2, MICU3 is exclusively expressed in the brain and enhances MCU activity in non-neuronal cells. Our studies are, to our knowledge, the first to uncover the physiological significance of the neuronal-specific expression of MICU3. We reason that the MICU3-mediated tuning of mitochondrial Ca^2+^ uptake is critical for the coordination of ATP synthesis and consumption in firing nerve terminals. However, mitochondrial Ca^2+^ uptake likely conveys risks as well, as overloading Ca^2+^ in the mitochondrial matrix is thought to facilitate the permeability transition of the inner membrane, in turn leading to necro-apoptotic cascades (Krieger and Duchen, 2002). Whether neuronal mitochondria have specific mechanisms at work to mitigate this risk is an important question that requires further investigation. Given the metabolic vulnerability of the nervous system, our identification of the molecular machinery for activity-dependent regulation of mitochondrial ATP, future studies should strive to determine whether mutations in the components of the MCU complex give rise to neurodegeneration in human patients.

It is interesting to note that in addition to glucose and lactate, the brain can also, under some conditions, be fueled by ketone bodies. How this fuel source is utilized and how its combustion is regulated is not completely understood (Lutas and Yellen, 2013). Given that strict ketogenic diets are used to curtail certain forms of intractable epilepsies (Neal et al., 2008), it will be interesting in the future to dissect how the regulation of ketone usage differs from other fuel sources at the synapse, and what is the physiological consequence of this differential regulation on synaptic function and cognitive performance. The metabolic plasticity of the brain also explains why genetic or pharmacological manipulation of metabolic pathways sometimes fail to exhibit dramatic cognitive phenotypes. Despite its crucial role in the regulation of presynaptic glycolysis, knockout models of GLUT4 survive (Abel et al., 1999) and have not been reported to have gross cognitive impairment, perhaps due to compensatory mechanisms. Similarly, inhibition of pyruvate import into neuronal mitochondria does not impair gross ATP production or survival, and merely results in metabolic reprogramming (Divakaruni et al., 2017). Such robust metabolic plasticity in the mammalian brain most likely serves to ensure that cognitive performance is faithfully maintained in the face of changing nutrient availability.

## Acknowledgements

We thank members of the Ryan lab for frequent discussions about this work and Adam Cohen (Harvard) for generously providing the spiking HEK cells.

## STAR METHODS

### Contact for Reagent and Resource Sharing

Further information and requests for resources and reagents should be directed to and will be fulfilled by the Lead Contact Timothy A. Ryan at.

### Experimental Model and Subject Details

#### Animals

All animal-related experiments were performed with wild-type rats of the Sprague-Dawley strain (Charles River code 400, RRID: RGD_734476.) in accordance with protocols approved by the Weill Cornell Medicine IACUC.

#### Primary Neuronal Culture

Hippocampal neurons were isolated from 1-to 2-day-old rats of mixed gender, plated on poly-ornithine-coated coverslips, transfected 7 days after plating, and imaged 14-21 days after plating as previously described (Ryan, 1999). Neurons were maintained in culture media composed of MEM (Thermofisher Scientific S1200038), 0.6% glucose, 0.1 gm/l bovine transferrin (Millipore 616420), 0.25 gm/l insulin, 0.3 gm/l glutamine, 5% fetal bovine serum (Atlanta Biologicals S11510), 2% B-27 (Thermofisher Scientific 17504-044), and 4 μm cytosine β-d-arabinofuranoside. Cultures were incubated at 37°C in a 95% air/5% CO2 humidified incubator for 14–21 days prior to use.

### Method Details

#### Plasmid Constructs

The following previously published DNA constructs were used: vGLUT1-pHluorin (Voglmaier et al., 2006), Syn-ATP (Rangaraju et al., 2014), ER-GCaMP6-150 and ER-GCaMP6-210 (de Juan-Sanz et al., 2017), α2 δ1 and β1B (Yan et al., 2013), CaV 2.1 (Hoppa et al., 2012), Ace-mNeon (Gong et al., 2015), MCU-v5-HIS (Baughman et al., 2011) (addgene 31731) was cloned into the BamH1 and Xho1 sites of the lentiviral expression vector pLenti-MP2 (Enomoto et al., 2013) (addgene 36097). We designed Mito^4x^-GCaMP6f to express 4 consecutive copies of the signal peptide of COX8 (MSVLTPLLLRGLTGSARRLPVPRAKIHSLGDP) and a short linker (RSGSAKDPT) before the sequence of GCaMP6f (Chen et al Nature 2013), because frequent mislocalization in the cytosol was observed when neurons expressed GCaMPs targeted by a single copy of the signal peptide (Biophysical Journal Cohen Lab 1x-mitogcamp). To target the pH-sensitive flurophore pHluorin to the mitochondrial matrix (Mito^4x^-pHluorin), we replaced GCaMP6f in Mito^4x^-GCaMP6f for pHluorin using AfeI and AgeI sites. For targeting GCaMP6f and jRCaMP1b to the outer membrane of mitochondria, we designed constructs to express the indicators after the first 33 amino acids of human TOM20 (MVGRNSAIAAGVCGALFIGYCIYFDRKRRSDPN) (Kanaji et al JCB 2000) and a short linker (TGS). To generate a vector to express both Mito^4x^-GCaMP6f and OMM-jRCaMP, we generated by gene synthesis a construct to express under the CMV promoter OMM-jRCaMP1b followed by a short linker (TG), the P2A sequence (ATNFSLLKQAGDVEENPGP) and a short linker (GSTA) that introduces a BamHI site (GGATCC). We subsequently cloned Mito^4x^-GCaMP6f between BamHI and EcoRI sites. We confirmed that this vector successfully expressed and maintained localization of both indicators in HEK cells, but it did not consistently do so in primary neurons as OMM-jRCaMP1b appeared frequently mislocalized in the cytosol despite proper localization of Mito^4x^-GCaMP6f to mitochondrial matrix.

To generate an shRNA resistant variant of rat MICU3, we codon-optimized rat MICU3 sequence and introduced at least 5 silent mutations in the regions complementary to shRNA designed (for shRNA A: shRNA resistant MICU3 region GCAATGAAATGGTCGACAA contains 5 mismatches; for shRNA B: shRNA resistant MICU3 AGAGCTGCACAGTCGTTAA contains 7 mismatches). This construct was used for rescue experiments (Fig S3) and overexpression of MICU3 in spiking HEK cells (Figure 5).

All constructs generated by gene synthesis (GeneArt Gene Synthesis, ThermoFisher) were codon-optimized for mouse expression using the GeneOptimizer tool (ThermoFisher)

We used the pLKO.1 vector (addgene 10878) for expression of shRNAs against rat MCU which was obtained from Origene Technologies (TR704979B, GCTACCTTCTCGGCGAGAACGCTGCCAGTT) and rat MICU3 (shRNA A: GAAACGAGATGGTGGATAA; shRNA B: GGAACTTCATAGCAGATAA).

### Lentiviral Production and Application

HEK 293FT cells were transfected by calcium phosphate with lentiviral constructs along with the associated packaging plasmids psPAX2 (a gift of Didier Trono, Addgene 12260) and pMD2.G (a gift of Didier Trono, Addgene 12259). 16 hours post transfection, media was changed to serum free viral production media: Ultraculture (Lonza), 1% (v/v) Penicillin-Streptomycin/L-glutamine, 1% (v/v) 100 mM sodium pyruvate, 1% (v/v) 7.5% sodium bicarbonate and 5 mM sodium butyrate. HEK 293FT supernatants were collected at 46 hours post transfection and filtered through a 0.45 μm cellulose acetate filter. Viral supernatants were then concentrated and buffer exchanged into neuronal culture media (MEM, 5% FBS, 2 μM ARA-C, 2% B-27) using an Amicon Ultra-15 100K MW Cutoff Centrifugal Filter (Millipore). Samples were aliquoted and stored at −80°C until use.

For imaging experiments, 1 μL of virus was added to neurons 3-4 days in vitro in a 6 mm cloning cylinder. For qPCR, 5 μL were added per well of a 24 well plate and in both cases after 3 days of addition, media was replaced with fresh media. Due to the long half-life of MCU (Cohen LD PLoS One 2013 and Seok Heo PNAS 2018), all experiments were performed at least 10 days after viral transduction to ensure protein knockdown.

### Live Imaging of Neurons

Imaging experiments were performed on a custom-built laser illuminated epifluorescence microscope with an Andor iXon+ camera (model #DU-897E-BV). TTL-controlled Coherent OBIS 488 nm and 561 nm lasers were used for illumination. Images were acquired through a 40X 1.3 NA Fluar Zeiss objective. Coverslips were mounted in a laminar flow perfusion chamber and perfused with Tyrodes buffer containing (in mM) 119 NaCl, 2.5 KCl, 2 CaCl_2_, 2 MgCl_2_, 50 HEPES (pH 7.4), 5 glucose or 1.25 lactate and 1.25 pyruvate, supplemented with 10 µM 6-cyano-7nitroquinoxalibe-2, 3-dione (CNQX), and 50 µM D,L-2-amino-5phosphonovaleric acid (APV) (both from Sigma) to inhibit post-synaptic responses. Action potentials were evoked in neurons with 1 ms pulses creating field potentials of ~10 V/cm via platinum–iridium electrodes. Temperature was maintained at 37°C using a custom-built objective heater jacket in all experiments except for those shown on Figure 3, which were performed at 26°C to eliminate the previously-observed modulation of presynaptic Ca^2+^ entry triggered by CPA at 37°C (de Juan-Sanz et al 2017; Emptage et al Neuron 2001; Liang et al 2002 J of Neurophysiology). Luminescence imaging of the presynaptic ATP reporter, Syn-ATP, was performed as previously described (Rangaraju et al., 2014). Although all our ATP measurements are pH-corrected as previously described (Rangaraju et al., 2014), given that we did not observe significant differences in pH changes (data not shown) when comparing wild type neurons and MCU/MICU3 KD neurons, here we do not propagate errors of pH measurements into the final error shown in ATP measurements.

NH_4_Cl solution for alkalization of pHluorin-containing vesicles had a similar composition as Tyrodes buffer except it contained (in mM): 50 NH_4_Cl and 69 NaCl. Ionomycin (Alomone Labs) at 500 µM final concentration was applied in pH 6.9 Tyrodes buffer containing 4mM CaCl_2_. Neurons were incubated for 5 minutes with 2 µM oligomycin (Enzo Life Sciences) in Tyrodes buffer to block mitochondrial ATP production.

### Culture and transfection of Spiking HEK293 cells

Spiking HEK cells stably expressing the voltage-gated sodium channel Na_V_ 1.3 and the inward rectifying potassium channel K_IR_ 2.1 were obtained from Dr. Adam Cohen (Harvard University) (Park et al., 2013). K_IR_ 2.1 expression is controlled by a tetracycline Tet-On system. To induce a consistent expression of K_IR_ 2.1, spiking HEK cells were cultured in the presence of 5 μg/ml doxycycline for 3-4 days before imaging. Spiking HEK cells were plated in glass coverslips and transfected at a 20% confluency using established calcium phosphate protocols (Kingston et al., 2001), adding a total amount of 7 μg of DNA per 35mm culture plate. For Ca^2+^ signaling experiments, we co-transfected 3 separate plasmids expressing CaV2.1, α2 δ1 and β1B together with the indicator of interest, conserving a ratio of DNA amounts of 1.5 / 1 / 1 / 1 respectively. When spiking HEK cells were additionally co-transfected with MICU3, the amount of this plasmid was 2-fold compared to α2 δ1, β1B or the indicator of interest. Cells were imaged 48h to 72h after transfection.

### Live Imaging of Spiking HEK293 cells

Imaging experiments were performed in the same custom-built setup used for neurons. Spiking HEK cells were imaged at a confluency of ~75%, which we found prevents their known spontaneous firing behavior induced at confluency by intercellular electrical coupling through gap junctions (Park et al., 2013). Single action potentials were evoked on-demand in Spiking HEK cells with a 10 ms pulse, creating a field potential of ~10 V/cm via platinum–iridium electrodes. Cells were imaged in the same Tyrodes buffer as neurons, except we removed CNQX and APV and varied Ca^2+^ concentrations as indicated, ranging from 0.5mM to 4mM with the corresponding change in MgCl_2_ concentrations to maintain osmolality in each buffer. Transfected areas of the dish were randomly chosen and typically contain 5 to 15 transfected cells. To image voltage changes in the HEK plasma membrane using AcemNeon (Fig. S2) (Gong et al., 2015), images were collected at 100Hz. For Ca^2+^ signaling experiments cells were imaged at 1Hz. We confirmed that Spiking HEK cells Ca^2+^ responses induced by field stimulation are stable for at least 30 minutes (not shown). When imaging two different Ca^2+^ indicators in the same HEK cell (Figure 4), we first imaged jRCaMP1b-based indicators, waited 2min and imaged GCaMP-based indicators in the same experimental conditions. The power of the 488 nm laser at the back aperture was never more than 0.3mW, as we have found that jRCaMP1b becomes unresponsive after strong 488 illumination. Whereas Mito^4x^-GCaMP6f appeared well localized in HEK mitochondria, we observed OMM-jRCaMP1b mislocalization in certain cases. HEK cells where the OMM-jRCaMP1b pattern did not match the Mito^4x^-GCaMP6f were excluded from the analysis.

### RNA Isolation and Quantitative PCR

Total RNA was isolated from primary dissociated cortical neuron cultures using TRIzol (Thermo Fisher) and chloroform (Sigma) along with RNeasy Mini Kits (Qiagen). RNA was reverse transcribed using SuperScript IV VILO Master Mix with ezDNase Enzyme using manufacturer recommended protocol. cDNA was used in qPCR reactions containing TaqMan Fast Advanced Master Mix (Thermo Fisher) and TaqMan Gene Expression Assays (Thermo Fisher). Relative mRNA expression was determined by normalizing to *Actb* using the ΔΔC_t_ method. Specific TaqMan assay information can be found in the Key Resources Table.

### Quantification and Statistical Analysis

#### Image Analysis and Statistics

Images were analyzed using the Image J plugin Time Series Analyzer where ~10-20 regions of interest (ROIs) of ~2 µm corresponding to responding synaptic boutons were selected and the fluorescence was measured over time. All fitting was done with OriginPro v8 as previously described (Balaji and Ryan, 2007). Statistical analysis was performed with OriginPro v8 and GraphPad Prism v6.0 for windows. In most experiments, the Mann–Whitney U test was used to determine the significance of the difference between two conditions. For paired comparison of responses, the Kolmogorov-Smirnov test was used. P < 0.05 was considered significant and denoted with a single asterisk, whereas P < 0.01, P<0.001 and P < 0.0001 are denoted with two, three, and four asterisks, respectively. The n value, indicated in the figure legends for each experiment, represents the number of cells imaged.

#### Quantification of Synaptic Vesicle Fractional Retrieval Block

Endocytic time constants were calculated as previously described by fitting the fluorescent change after the stimulus to a single exponential decay (Armbruster and Ryan, 2011). The fractional retrieval block in endocytosis of vGLUT1-pH was calculated as the fraction of ΔF remaining after 2 times the average endocytic time constant of the control (2Ƭ) to maximum ΔF at the end of stimulation (ΔF_2τ_/ ΔF_max_).

In the case of oligomycin treatment (Fig 1C-E), individual (not average) endocytic time constants before treatment was used for each cell.

#### Analysis of Spiking HEK293 Cells

Dynamic responses were analyzed using the Image J plugin Time Series Analyzer, placing manually-drawn single regions of interest (ROIs) per cell. Fluorescence signals in response to electrical activity (ΔF) were normalized to the resting fluorescence (F_0_). The F_0_ value was additionally corrected for background autofluorescence measured in a nearby non-transfected region. For AcemNeon experiments, we averaged 10-15 single-stimulation trials to reduce signal-to-noise in our measurements. For Ca^2+^ signaling experiments, data was obtained from single trials and we applied inclusion and exclusion criteria with the following elements: 1) To avoid overestimating ΔF/F_0_ which would arise in cells with low F_0_ values, we set an arbitrary threshold such that F_0_/background > 1.25 to be included for further analysis. 2) As spiking HEK cells may present spontaneous responses (Park et al., 2013), we include cells that are silent before stimulation by selecting cells whose standard deviation of their baseline fluorescence is low, excluding any cells with a standard deviation of the ΔF/F_0_ before stimulation higher than 0.075. 3) Lastly, we only include cells with a response of OMM-jRCaMP1b 3 times higher than the standard deviation of the baseline fluorescence, allowing us to only use cell with reliable outer membrane responses. Criteria #3 is not applied to Mito^4x^-GCaMP6f responses as arbitrarily small responses must be included to determine the responsive of the matrix signal for a given OMM signal.

#### Mitochondrial Ca2+ measurements using Mito^4x^-GCaMP6f

For mitochondrial Ca^2+^ signaling experiments, data was obtained from single trials imaging at 5Hz (Figure 2, 4 and 5) or averaging 3 trials (Figure 3). We found that in some cases a small fraction of Mito^4x^-GCaMP6f appeared mislocalized in the cytosol, which would contaminate the quantification of peak mitochondrial responses. We leveraged the kinetic differences of cytosolic and mitochondrial Ca^2+^ responses to quantify exclusively mitochondrial Ca^2+^. Since mitochondrial responses persist much longer than those in the cytosol chose a 1 sec delay after the stimulus as an measure or the matrix response that is not corrupted by any contribution from possibly mislocalized probe. To do so, we averaged 5 frames of Mito^4x^-GCaMP6f fluorescence 1s after stimulation when cytosolic Ca^2+^ has returned to baseline, and we use this value through the text as the peak mitochondrial Ca^2+^ response. For clarity, in figures, we show responses that did not present mislocalization. To avoid overestimating ΔF/F_0_ in cells with low F_0_ values, we set an arbitrary threshold applied to all Mito^4x^-GCaMP6f experiments such that F_0_/background > 1.25 to be included for further analysis.

#### Resting mitochondrial Ca^2+^ estimates using Mito^4x^-GCaMP6f

This method relies on the experimental measurement Mito^4x^-GCaMP6f fluorescence at saturating [Ca^2+^] in mitochondria (F_max_), which is obtained by applying Tyrode’s solution containing 500µM ionomycin, 4mM CaCl_2_ and 0mM MgCl_2_ at pH 6.9 buffered with 25mM HEPES. Knowing the in vitro characteristics of GCaMP6f (Chen et al., 2013), baseline mitochondrial [Ca^2+^] (Ca_r_) is calculated from F_max_ using the following equation:

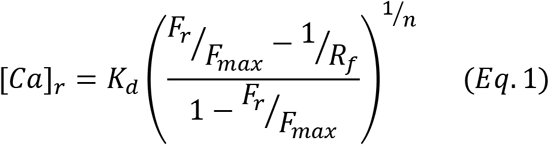

K_d_ is the affinity constant of the indicator, F_r_ is the measured fluorescence at rest, R_f_ is the dynamic range (Fsat/Fapo) and *n* is the Hill coefficient. Ionomycin application does not produces a change in mitochondrial matrix pH (data not shown; initial pH 7.203; ionomyicin-treated mitochondria pH 7.206). Neurons with apparent cytosolic mislocalization of Mito^4x^-GCaMP6f were excluded from these estimates.

#### Mitochondrial pH measurements

Mitochondrial pH measurements were made using Mito^4x^-pHluorin and axonal mitochondria were imaged. Neurons were briefly perfused with a Tyrode’s solution containing 100mM NH4Cl buffered at pH 7.4 25mM HEPES, which equilibrated mitochondrial pH to 7.4. The fluorescence value when pH stabilizes at 7.4 allows estimating resting mitochondrial pH value using the modified Henderson-Hasselbalch equation:

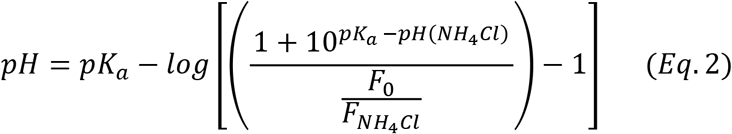

pKa is the pKa of pHluorin, 7.1, pH (NH_4_Cl) is the pH of 100mM NH_4_Cl buffer used, F_0_ is the fluorescence of Mito^4x^-pHluorin measured before NH_4_Cl perfusion, F_NH4Cl_ is the fluorescence of Mito^4x^-pHluorin measured upon NH_4_Cl perfusion when signal is stable.

