## supplementary figures and legends for "Molecular tuning of the axonal mitochondrial Ca^2+^ uniporter ensures metabolic flexibility of neurotransmission"

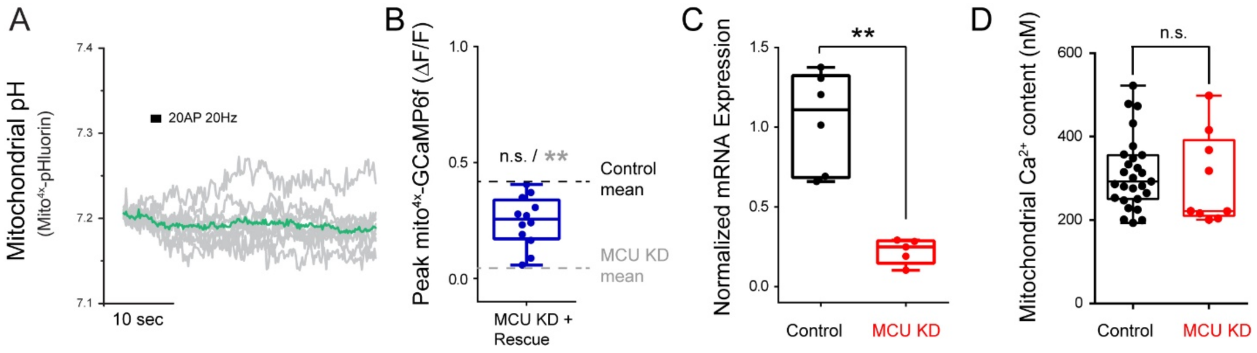


**Figure S1 (related to Figure 2). Physiological mitochondrial matrix parameters, validation of MCU KD in neurons with phenotypic rescue and quantitative PCR.** (A) Mitochondrial matrix pH does not change in response to electrical stimulation (black bar). Shown are individual (grey) and average (green) traces of Mito$^{4x}$-pHluorin, a matrix-targeted pHluorin construct. n = 10. (B) The reduction in mitochondrial $Ca^{2+}$ uptake in MCU knockdown neurons, measured with Mito$^{4x}$-GCaMP6f, is partially rescued with re-expression of shRNA resistant MCU-v5. (C) Relative mRNA expression levels of *MCU* mRNA, normalized to *actin*, in control and MCU KD cortical neurons. n= 5-6 samples. (D) Basal mitochondrial $Ca^{2+}$ content is not different in control and MCU KD neurons. n= 9-27 cells. Mean (nM); control, 312; MCU KD, 296. Error bars are SEM. n.s. not significant. p>0.05, **p<0.01.

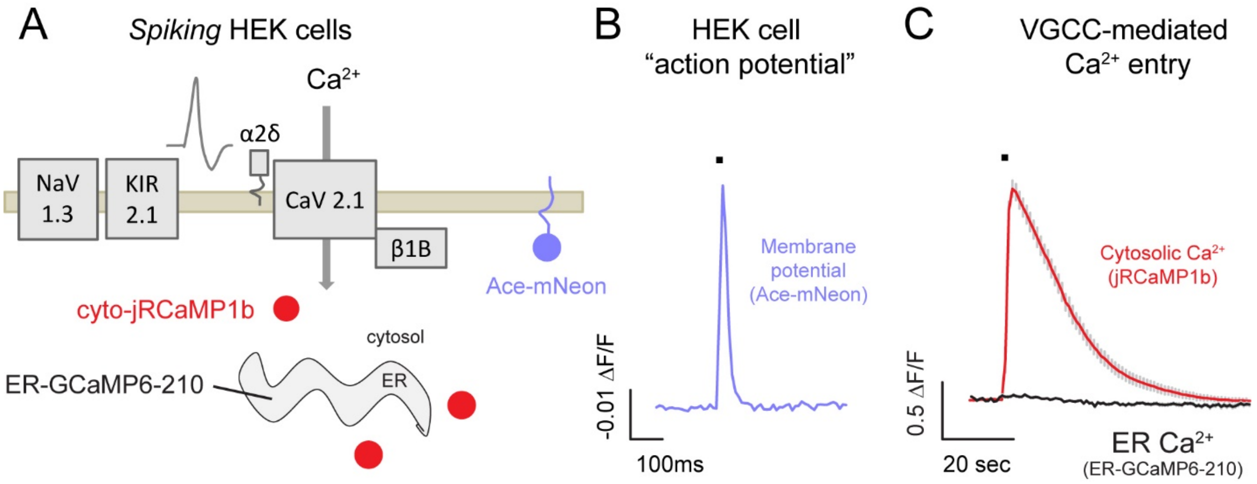


**Figure S2 (related to Figure 4). Characterization of membrane potential and $Ca^{2+}$ dynamics in spiking HEK cells.** (A) Schematic diagram of spiking HEK cells expressing voltage-activated $Na^+$ (NaV1.3) and $K^+$ (KIR2.1) channels, the subunits of voltage gated $Ca^{2+}$ channels (CaV2.1, α2 δ1, and β1b), and the membrane bound voltage sensor Ace-mNeon. (B) Average trace of Ace-mNeon in spiking HEK cells showing membrane depolarization in response to brief electrical field stimulation. n=7 cells. (C) Average traces of cytosolic and ER $Ca^{2+}$ fluxes in spiking HEK cells measured with jRCaMP1b and ER-GCaMP6-210 in response to stimulation. n= 26 cells. Error bars are SEM.

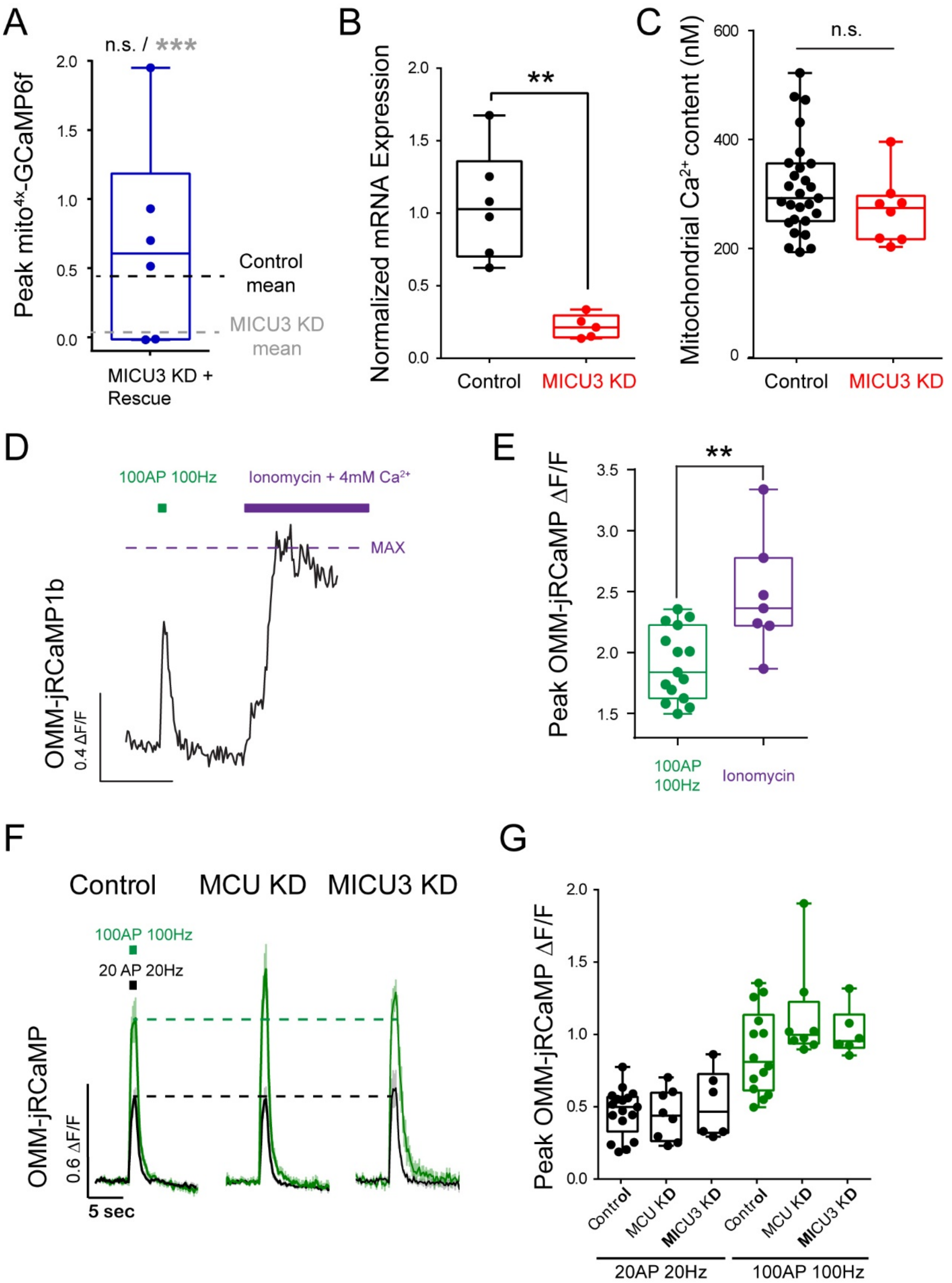


**Figure S3 (related to Figure 5). Functional rescue, validation of MICU3 KD and impact of MICU3KD on resting mitochondrial Ca2+ levels in neurons; verification that cytosolic Impact of MCU and MICU3 KD on cytosolic $Ca^{2+}$ levels responses to stimulation .** (A) The reduction in mitochondrial $Ca^{2+}$ uptake in MICU3 knockdown neurons, measured with Mito$^{4x}$-GCaMP6f, is fully rescued with re-expression of shRNA resistant MICU3. (B) Relative mRNA expression levels of *MICU*3, normalized to *actin*, in control and MICU3 KD cortical neurons. n=

5-6 samples. (C) $Ca^{2+}$ concentration in mitochondrial matrix is not changed in control and MICU3 KD neurons. n= 8-27 cells. Mean (nM); control, 311; MICU3 KD, 271. (D-E) Sample trace (D) and peak quantification (E) of a control neuron expressing the OMM-targeted $Ca^{2+}$ indicator jRCaMP1b in response to 100 AP, 100 Hz stimulation and ionomycin treatment to saturate the indicator showing that the indicator is not saturated during stimulation. n= 7-15 cells. Mean (ΔF/F); 100 AP 100Hz, 1.9; ionomycin, 2.5. (F-G) Average traces (F) and peak quantification (G) of OMM-jRCaMP1b to 20 AP, 20 Hz or 100 AP, 100 Hz stimulation in control, MCU KD and MICU3 KD neurons indicate that mitochondrial $Ca^{2+}$ uptake does not contribute to cytosolic $Ca^{2+}$ buffering. Mean (ΔF/F); 20AP 20 Hz, control, 0.47; MCU KD, 0.44; MICU3 KD, 0.52; 100 AP 100Hz, control, 0.88; MCU KD, 1.1; MICU3 KD, 1.0. n= 6-17 cells. Error bars are SEM.
